# Von Willebrand Factor Synergizes with Tumor-Derived Extracellular Vesicles to Promote Gastric Cancer Metastasis

**DOI:** 10.1101/2023.08.25.554906

**Authors:** Chen-yu Wang, Min Wang, Wei Cai, Chan-yuan Zhao, Quan Zhou, Xiao-yu Zhang, Feng-xia Wang, Chen-li Zhang, Yun Dang, Ai-jun Yang, Jing-fei Dong, Min Li

**Author notes:** These authors contributed equally (Chen-yu Wang, Min Wang). Correspondence: Min Li, Institute of Pathology, School of Basic Medical Sciences, Lanzhou University, Lanzhou, China, or Jing-fei Dong, Bloodworks Research Institute, Seattle, Washington, USA.

## Abstract

Cells of gastric cancer invade local tissue extensively and also metastasize through the circulation to remote organs. Patients with metastasized gastric cancer have poor outcomes. Cancer cells are known to release extracellular vesicles (EVs) that contribute to cancer progression, but how cancer cell-derived EVs promote cancer growth and metastasis remains poorly understood. We have recently reported that levels of circulating gastric cancer cell-derived EVs (gcEVs) and the adhesive ligand von Willebrand factor (VWF) are associated with cancer metastasis and poor prognosis of patients, but the underlying mechanism of this gcEV-VWF interaction was not known. Here we report results from a study designed to investigate the synergistic action of VWF and gcEVs in vitro and in mouse models. We showed that VWF in cancer-bearing mice was hyperadhesive and became microvesicle-bound. EV-bound VWF mediated the adhesion of gcEVs to the endothelium to disrupt endothelial integrity and facilitate the transendothelial migration of cancer cells and pulmonary metastasis. Reducing VWF adhesive activity by the metalloprotease ADAMTS-13 or promoting gcEV clearance by the scavenging factor lactadherin prevented pulmonary metastasis in mice. These results highlight the synergistic action of gcEVs and VWF in promoting gastric cancer metastasis and identifying new targets for its prevention.

**Key point: Author contributions:**

1. Hyperadhesive VWF becomes microvesicle-bound to induce endothelial leakage and promote the pulmonary metastasis of gastric cancer in mice.
2. Reducing VWF activity by ADAMTS-13 and accelerating microvesicle clearance by lactadherin reduces pulmonary metastasis of gastric cancer.

## INTRODUCTION

Gastric cancer (GC) is the third leading cause of cancer mortality worldwide, accounting for more than 800,000 deaths every year.^1-3^ The high mortality of GC is attributed to the direct invasion of cancer cells into adjacent organs and to metastasis through the circulation to remote organs. The hematogenous metastasis of GC is a multistep process during which cancer cells are released into the circulation through the poorly developed and “leaky” vessels of cancer tissue, adhere to the endothelium of remote organs, induce endothelial permeability in targeted organs, and then transmigrate through the endothelium to form new cancer loci.^4-6^ The critical questions are how circulating cancer cells selectively adhere to and transmigrate through the endothelium of a specific organ, which is fully developed and has limited permeability, and what molecules mediate the interaction between cancer cells and the endothelium.

Tumor-derived extracellular vesicles (EVs) have been shown to enhance the proliferation, invasion, and metastasis of cancers.^7-11^ Our studies further show that EVs from GC and adhesive ligand VWF may play a synergistic role in cancer metastasis.^12,13^ EVs are membrane-enclosed nanoscale particles derived from membrane fragments, intracellular granules, and secreted exosomes that are released by cells undergoing apoptosis or active microvesiculation.^14,15^ EVs are highly heterogeneous in size, structure, and cargo molecules.^16^ We have shown that (1) gastric cancer-derived EVs (gcEVs) bind and are endocytosed by endothelial cells to induce cytoskeletal movements and permeability, thus promoting the transendothelial migration of GCs^12^ and (2) GC synthesizes and releases the adhesive ligand VWF, which promotes GC-platelet interaction and metastasis.^17^ These studies led us to hypothesize that hyperadhesive VWF serves as a coupling factor to capture gcEVs to the endothelial cells of specific vascular beds, where gcEV-carried factors induce endothelial permeability and promote GC metastasis.^12^ This hypothesis was also developed because VWF multimers are synthesized in endothelial cells,^18^ through which cancer cells metastasize. Upon release, ultra-large and hyperadhesive VWF multimers are anchored to endothelial cells, elongated by the shear stress of blood flow, and released by the metalloprotease ADAMTS-13 (a disintegrin and metalloprotease with thrombospondin type motifs type 13), which cleaves the peptide Tye^165^-Met^166^ bond in the A2 domain.^19-21^ Although VWF primarily functions to initiate hemostasis, it also promote thrombosis and inflammation by facilitating the interaction between endothelial and blood cells.^22,23^ It is at this blood-endothelial interface that VWF has been increasingly recognized for regulating cancer growth and contributing to outcomes of cancer patients.^24-26^ However, the question remains as whether VWF serves as a marker of endothelial activation/injury or plays a causal role in cancer metastasis. Here, we discuss results from the study designed to investigate the synergistic activity of gcEVs and VWF in promoting GC metastasis and the means for mitigating this cancer-promoting activity.

## MATERIALS AND METHODS

### Mouse model

Mouse experiments were approved by the Institutional Animal Care and Use Committee of Lanzhou University. They were performed on adult 615 mice (a derivative strain from C57BL/6J mice) of 6-8 weeks old of both genders (Institute of Hematology, CAMS, Tianjin, China).

We used an adaptive transfer model to evaluate the pulmonary metastasis of GC through the circulation, as we and others previously described.^12,27^ We discussed the rationale for using this model in the Supplemental Methods section. Briefly, cells from the mouse forestomach carcinoma line (MFC, CCTCC, Shanghai, China) were cultured in RPMI-1640 medium supplemented with 10% FBS until confluence. They were then detached with 0.5% EDTA, washed, and resuspended in PBS. Anesthetized mice received a single bolus infusion of MFC cells (1.5×10^6^ in 200 µl) through the tail vein. The MFC-infused mice received every three days for 14 days of (1) gcEVs generated from MFC (Supplemental Methods); (2) VWF-bound gcEVs (VWF^+^gcEVs) obtained by incubation of gcEVs with 10 µg of recombinant (r) human VWF (Sino Biological, Beijing, China); (3) gcEVs and 5μg of lactadherin (Signalway Antibody, USA), (4) VWF^+^gcEVs and 5 μg of lactadherin; (5) VWF^+^gcEVs and 5 μg of human rADAMTS-13 (MyBioSource, USA); and (6) VWF^+^gcEVs with rADAMTS-13 and lactadherin, each at 5 μg/mouse. Control mice received an equal volume of PBS, 10 µg/kg of rVWF, 5 μg of lactadherin, or 5 μg of rADAMTS-13 in the absence of gcEVs and MFC cells. The rVWF used in this study was not cleaved by ADAMTS-13 and thus hyperadhesive (**Supplemental Figure S1**). The rADAMTS-13 used for treatment cleaved both human and mouse VWF when exposed to shear stress,^28^ achieving 72% and 65% cleavage measured by mass spectrometry assay.^29^ Blood samples were collected through cardiac puncture immediately after euthanasia (anticoagulant: 0.32% sodium citrate), and the lungs were also collected and evaluated for pulmonary metastasis.

### EV isolation and immune electron microscopy

Blood was centrifuged at 1,000×g for 5 min and then at 10,000×g for 10 min to remove cells and large aggregates. The supernatant was centrifuged at 100,000×g for 60 min at 4°C twice to collect gcEVs pellets, which were processed for immunoelectron microscopy.^30^ Briefly, EVs were fixed with 4% paraformaldehyde (Solarbio, Beijing, China), washed, and blocked with 1% albumin. They were then incubated with a rabbit anti-VWF antibody (Abcam, Cambridge, MA) followed by incubation with an immunogold-coupled goat-anti-rabbit IgG.

### VWF measurements

VWF antigen (VWF:Ag) was measured using ELISA (Shanghai Jianglai Biotechnology, China). VWF adhesive activity was measured by collagen binding (VWF:CB, Supplemental Methods), as we previously described.^31,32^

### Characterization of EVs using flow cytometry

Blood samples were centrifuged at 150×g for 20 min at RT to obtain platelet-rich plasma (PRP), which was centrifuged at 1,500×g for 20 min at RT to collect platelet-poor plasma (PPP). The PPP was incubated with detecting agents to identify EVs using flow cytometry (BD FACSAria™ Ⅱ, Becton Dickinson, San Jose, CA), as we have previously described.^12^ The flow cytometry method was discussed in details in the Supplemental Methods section, including how gcEVs were defined.

### Vascular permeability

Endothelial permeability was measured using two systems to cross-validate data: transendothelial electrical resistance and a transwell assay, as we previously reported with minor modifications.^12,33^ For the transwell assay, human umbilical cord endothelial cells (HUVECs; CCTCC) were plated in collagen coated transwells and cultured in DMEM/F12 medium (HyClone) containing 10% FBS (Gibco) until confluence. They were incubated with gcEVs, VWF^+^gcEVs, or VWF for 8hrs at 37°C, followed by incubation with fluorescein isothiocyanate (FITC)-Dextran (100 mg/mL, Sigma-Aldrich) for 60 min at 37°C. FITC intensity was measured in the medium from the bottom wells using a plate-reader (Molecular Devices, San Jose, CA).

### Assay for gcEV-induced activation of endothelial cells

Confluent HUVECs on gelatin-coated plates were washed with PBS and treated with gcEVs for 30 min at 37°C and then examined for releasing VWF and surface adherent gcEVs. Cells treated with 25 μM of histamine (Sigma-Aldrich)^34^ served as positive controls. For VWF release, the conditioned media from treated HUVECs were centrifuged at 3000×g for 5 min and the supernatant was analyzed for VWF:Ag using ELISA (Sino Biological Inc.). For gcEV binding, HUVECs on gelatin-coated plates were incubated with gcEVs labeled with the fluorescent dye PKH-26 (Sigma-Aldrich, 10 min at RT). After washing, the cells were processed for immunofluorescence staining.

### Immunofluorescence staining of endothelial VWF

HUVECs were fixed with 4% paraformaldehyde (Solarbio, Beijing, China) for 20 min, washed, permeabilized with PBS-T (0.1% Tween-20), and blocked with 1% BSA for 1 hr at RT. They were then stained with a rabbit anti-VWF antibody (1:50, Abcam) overnight at 4°C and viewed under a Zeiss LSM900 confocal microscope (40×/1.4NA objective lens, Carl Zeiss, White Plains, NY).

### Cell adhesion and transmigration assays

For cell adhesion, HUVECs were plated at 1×10^5^ cells/ml onto 96-well plates and allowed to grow until confluence. They were then incubated with PKH-26-labeled gcEVs or VWF^+^gcEVs(2,000-3,000/µl) with or without the VWF-blocking antibody (Abcam) for 0.5 or 1 hr. After washing, treated endothelial cells were incubated for 30 min at 37°C with 1×10^4^/ml of cells from the two human gastric cancer lines AGS or HGC-27 (Cell Bank of the Shanghai Institute of Cell Biology, Chinese Academy of Sciences) that were pre-labeled with the membrane-permeable fluorescent dye carboxyfluorescein succinimidyl ester (CFSE; Bestbio Biotechnology, Shanghai, China). Upon diffusion into cells, CFSE binds covalently to lysine residues of proteins to stably emit green fluorescence (excitation: 488 nm and emission: 517 nm). After incubation, HUVECs were washed and CFSE^+^ cells adherent to endothelial cells were counted under a Nikon Eclipse Ti2-E inverted fluorescence microscope. HUVECs treated with 25 μM of histamine (Sigma-Aldrich) and serum-free medium served as positive and negative controls, respectively.

For transendothelial migration, HUVECs were plated at 1×10^5^ cells/ml into transwell inserts (pore size: 8 μm; Corning Inc., Somerville, MA) and allowed to grow until confluence. They were incubated with gcEVs or VWF^+^gcEVs, with or without the presence of the VWF-blocking antibody (Abcam). After washing, the cells were incubated with 5×10^4^/well of PKH26^+^ AGS or HGC-27 cells for 24 hrs at 37°C and the PKH26^+^ cells that migrated to the opposite side of the membrane were counted under a Nikon Eclipse TE2000-U fluorescence microscope.^35^

### Quantifying pulmonary metastasis of gastric cancer

Two quantitative measures were used to define the role of gcEVs in promoting hematogenous metastasis: total weight of the lungs and cancer load by histopathology. Both methods are widely used and well established in studying pulmonary metastasis of the lungs, including our own studies.^36-38^ Briefly, the lungs were collected from mice receiving various treatment regimens immediately after euthanasia and weighted. They were then fixed with 4% formaldehyde and sectioned. The tissue sections were stained with hematoxylin-eosin^39,40^ and scanned with the Pannoramic DESK system to acquire images using Pannoramic Case Viewer 2.4 (3DHISTECH, Hungary). The cancer load (%) was calculated as the cross-sectional area of cancer mass/the total area of the lungs×100.

### Bioinformatics analysis

We analyzed public databases to link VWF to ECs in cancer tissue. First, the Cancer Genome Atlas (TCGA) database was used to compare levels of VWF mRNA in stomach adenocarcinoma (STAD) and surrounding normal tissues (https://genome-cancer.ucsc.edu/). Second, we plotted VWF expression against different clinical stages of STAD from TCGA. Third, we analyzed the VWF expression for differential outcomes using the public GSE54129 and GSE19826 databases (https://www.ncbi.nlm.nih.gov/geo/). Finally, we used the “Immune-Gene” module of the Tumor Immune Estimation Resource (TIMER2; http://timer.cistrome.org/)^41^ to analyze the relationship between VWF expression and EC in STAD from TCGA. The gene expression was represented as log2 TPM. Cancer-associated ECs were analyzed using the xCELL, MCP-counter (Microenvironment Cell Populations-counter), and EPIC (Estimate the Proportion of Immune and Cancer cells) algorithms.^42-44^

### Statistical analyses

Quantitative data were reported as mean±SD and analyzed using either two-tailed t-test for comparison between two groups or analysis of variance (ANOVA) followed by post-hoc testing for multiple groups. The difference between groups was considered statistically significant with a *p* value of <0.05. Statistical analyses were performed using GraphPad Prism software 9.4.0. For bioinformatic analyses, the *p* values and partial correlation (COR) values were obtained using a purity-adjusted Spearman’s rank correlation test.

## Data Sharing Statement

The public database can be found through URL as listed in the article. For original data, please contact Dr. Min Li at.

## RESULTS

### GcEVs promoted cancer metastasis in mice

We established an adaptive transfer mouse model to study the role of EVs in the hematogenous metastasis of gastric cancer to the lungs, with treatment regimens shown in **Figure 1A**. All mice survived during the study period of 14 days. This model allowed us to study the hematogenous metastasis of cancer cells without the confounding influence of primary cancer mass, which varies significantly in size and causes systemic inflammation, which promotes metastasis.^45^ We found that MFC cells infused through the tail vein metastasized to the lungs to form new cancer loci that grew over time, as shown by metastatic nodules on the surface of the lungs and by histopathology (**Figure 1B**). Plasma levels of gcEVs, which were defined by the expression of CD326 and binding annexin V, increased significantly 14 days after MFC infusion (**Figure 1C**), accounting for more than 80% of the total PS-expressing EVs in the circulation (**Supplemental Figure S2**). When given together with MFC cells, gcEVs (**Figure 1C**) accelerated cancer metastasis to the lungs as detected by histopathology (**Figure 1D**) and quantified by the weight (**Figure 1E**) and total tumor area (**Figure 1F**) of the lungs. This metastasis-promoting effects of gcEVs was not caused by different activities of gcEVs in the plasma of MFC-implanted mice and those made in vitro because gcEVs obtained by the two methods were similar in size, CD326 expression, and PS exposure (**Supplemental Figure S3**). The metastasis-promoting effect of gcEVs was significantly reduced in mice receiving MFC cells and gcEVs together with 5µg of lactadherin (**Figure 1G**), which promotes the clearance of PS-expressing EVs.^12,30^ Mice receiving lactadherin also had lower levels of circulating PS^+^gcEVs (**Supplemental Figure S4**). These data suggest that gcEVs promoted the pulmonary metastasis of gastric cancer cells and this gcEV activity was reduced by enhancing the clearance of PS^+^ EVs.

**Figure 1:**
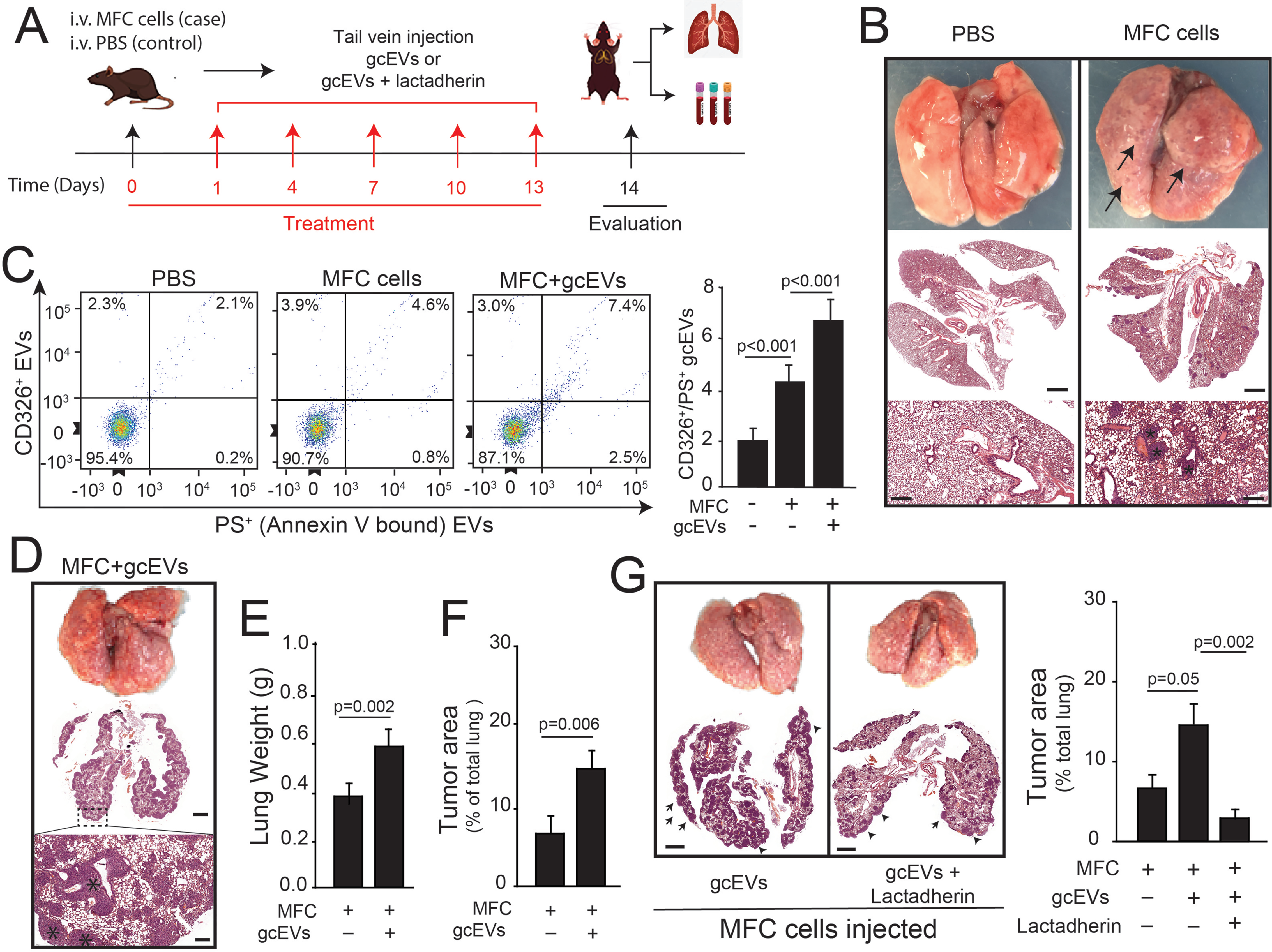
GcEVs promoted cancer metastasis in mice. (**A**) The experimental schedule for the mouse adaptive transfer model used in this study. Mice were infused with 1.5×10^6^/mouse of MFC cells alone or with additional factors. (**B**) Top panel: gross appearance of the lungs dissected from mice 14 days after being infused with MFC cells or an equal volume of the vehicle PBS, showing cancer nodules (darker colored spots indicated by arrows). Middle panel: H&E-stained lungs from the same mice (bar=1 mm). Bottom panel: enlarged images from the middle panel show dark brown metastasized masses (dark stars, bar=200 μm). The images are representative of 15 mice examined. (**C**) CD326^+^/PS^+^gcEVs in plasma of mice infused with MFC cells alone or with exogenous gcEVs (n = 6, paired *t*-test). (**D**) The lung metastasis from a mouse 14 days after infusion of MFC and gcEVs. Top: gross appearance. Middle and low panels (bars=1 mm and 200 μm, respectively): H&E-stained lungs from the mouse show dark brown metastatic loci, indicated by stars. The weights (**E**) and cancer-bearing areas (**F**) of the lungs from 6 treated mice per group (*t*-test). (**G**) Left: the metastasis nodules (top) and cancer-bearing areas (bottom, arrows: cancer loci) of the lungs from a mouse infused with MFC and gcEVs with and without lactadherin (bar=1 mm). Right: quantitative data from five independent experiments (one-way ANOVA).

### GcEVs activated ECs to release hyperadhesive VWF

Plasma samples from mice receiving MFC cells contained higher levels of VWF:Ag, which increased further in mice infused with MFC together with gcEVs (**Figure 2A**). VWF:CB was also higher in mice receiving MFC, and increased further in mice receiving MFC and gcEVs (**Figure 2B**). The VWF:CB-to-VWF:Ag ratio was significantly higher in mice receiving MFC cells and further increased in mice receiving MFC and gcEVs (**Figure 2C**), suggesting that VWF in these mice was intrinsically hyperadhesive. This notion was further supported by detecting increasing amounts of VWF on PS^+^EVs in mice receiving MFC and gcEVs compared to mice with MFC alone (**Figure 2D**). IEM further verified that VWF was EV-bound (**Supplemental Figure S5A**). Numbers of VWF^+^/PS^+^ EVs were closely associated with levels of VWF:Ag (R^2^ = 0.5175, p<0.001, **Supplemental Figure S5B**). Levels of both VWF^+^/PS^+^ EVs (**Figure 2E**) and total PS^+^ EVs (**Supplemental Figure S4)** were reduced in mice receiving lactadherin together with MFC cells and gcEVs. VWF:Ag (**Figure 2F**) and VWF:CB (**Figure 2G**) were also reduced in the lactadherin-treated mice. These data suggest that (1) gcEVs stimulated endothelial cells to release hyperadhesive VWF that bound to gcEVs and (2) lactadherin reduced levels of VWF:Ag and its adhesive activity (VWF:CB) by removing VWF-bound gcEVs.

**Figure 2:**
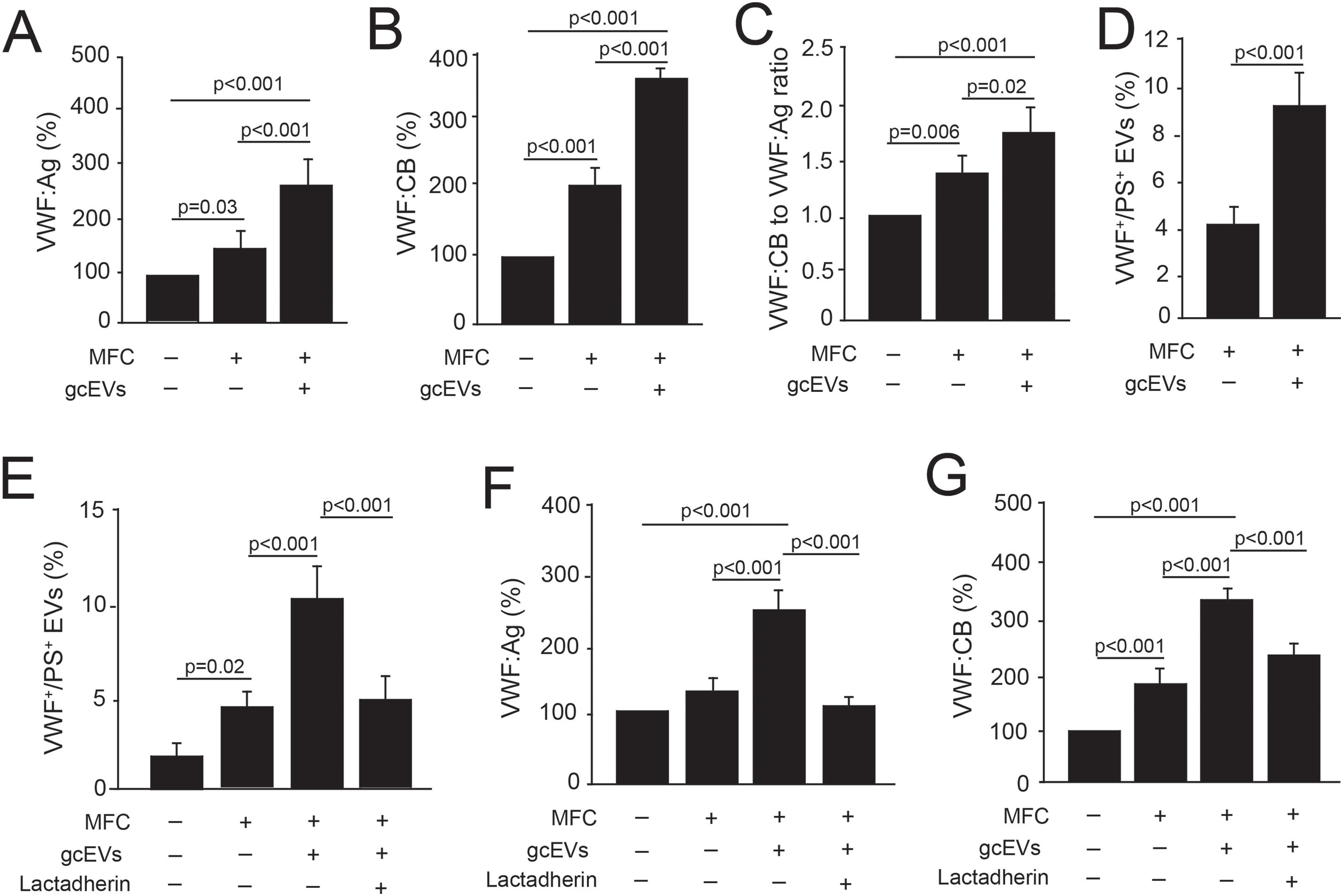
VWF in mice infused with MFC and gcEVs was hyperadhesive. Plasma VWF:Ag (**A**), VWF:CB (**B**), VWF:CB to VWF:Ag ratio (**C**), and levels VWF^+^/PS^+^ EVs (**D**) from mice infused with MFC cells with and without gcEVs (n = 5/group, one-way ANOVA). The percentage of VWF^+^/PS^+^ EVs (**E**), plasma VWF:Ag (**F**) and VWF:CB (**G**) in plasma of mice 14 days after infusion with MFC cells with and without gcEVs and lactadherin (n = 5/group, one-way ANOVA).

### VWF enhanced gcEV-induced cancer metastasis

The data in Figure 2 raised the question as to whether hyperadhesive VWF on gcEVs served as a bridge molecule linking gcEVs to endothelial cells, thus enhancing the effect of gcEVs on promoting endothelial permeability and the pulmonary metastasis of MFC cells. We investigated this possibility using the adaptive transfer model to answer two questions: (1) Does increasing the amount of hyperadhesive VWF further enhance the pulmonary metastasis of MFC cells, and (2) does reducing VWF adhesive activity by ADAMTS-13 reduce the pulmonary metastasis of MFC cells? We found that mice receiving MFC cells and gcEVs together with hyperadhesive rVWF (**Supplemental Figure S1**) had a greater rate of pulmonary metastasis of MFC cells (**Figure 3A-3C**). These mice had stiffer lungs, more visible metastatic nodules, and multiple metastasis loci detected by histopathology. ADAMTS-13 reduced pulmonary metastasis in mice receiving MFC and gcEVs with and without exogenous VWF (**Figure 3A-3C**). The protective effect of ADAMTS-13 was additive to that of lactadherin. ADAMTS-13-treated mice also had decreased levels of VWF^+^/PS^+^ EVs (**Figure 3D**), which served as a surrogate marker for VWF hyperadhesive activity. These results suggest that hyperadhesive VWF synergized with gcEVs to promote the pulmonary metastasis of circulating MFC cells. This metastasis-promoting activity was significantly enhanced by hyperadhesive VWF and reduced by increasing VWF cleavage by ADAMTS-13.

**Figure 3:**
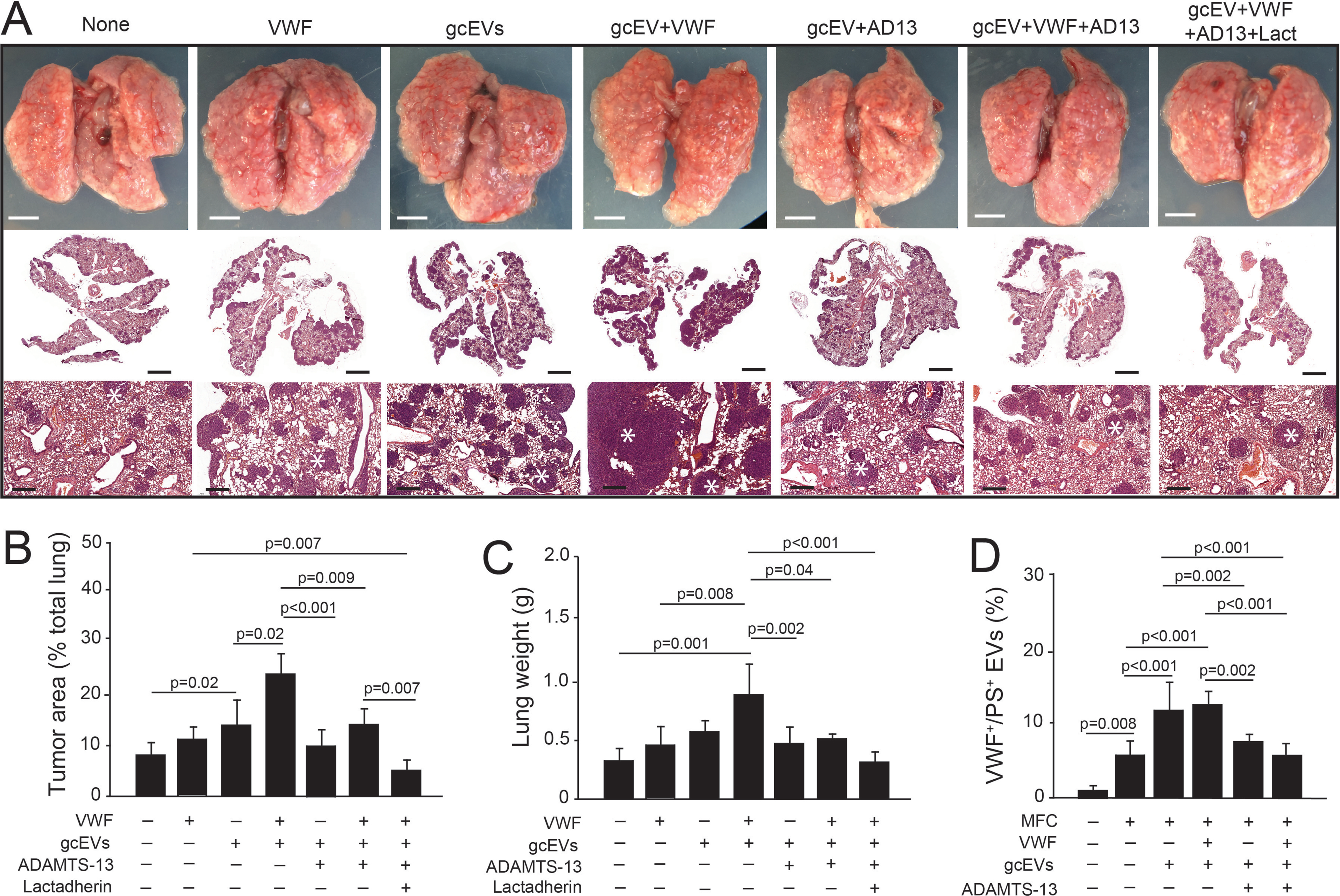
VWF enhanced gcEV-induced cancer metastasis. (**A**) The metastasis nodules on the surface of the lungs from mice 14 days after they were infused with MFC cells and other agents. Images are representative from 5-10 independent experiments (AD13: ADAMTS-13, lact: lactadherin). Top: gross appearance of the lungs with metastatic nodules. Middle and low: H&E-stained lungs at different magnifications (top bar=2.5 mm, middle bar=1 mm, bottom bar=200 μm) show dark brown metastatic cancer loci (white star at the bottom panels). Cancer-bearing areas (**B**) and weights of the lungs (**C**) were determined through quantitative analyses of the lungs from multiple mice (n = 5/group, one-way ANOVA). (**D**) Levels of VWF^+^/PS^+^ EVs in plasma samples of mice 14 days after infusion with MFC cells with and without gcEVs, VWF and ADAMTS-13 (n = 5/group, one-way ANOVA).

### GcEVs induced endothelial permeability in a VWF-dependent manner

We therefore examined the synergy among VWF, gcEVs, and endothelial cells in promoting the pulmonary metastasis of gastric cancer in more controlled in vitro systems to reduce the confounding influences existed in tumor-bearing mice (e.g., variable size of cancer masses). GcEVs released from cells of three human gastric cancer lines that were subjected to serum starvation (48 hrs at 37°C), which induced cells to undergo apoptosis to release gcEVs. GcEVs from the HGC-27, AGS, and BGC-823 cell lines of gastric cancer obtained using this method (Supplemental Methods) were similar in size and PS expression (**Supplemental Figure S6**). We found that human gcEVs from HGC-27 cells stimulated cultured HUVECs to release VWF (**Figure 4A**) and VWF-bound EVs (**Figure 4B**) in a dose-dependent manner. EVs from AGS and BGC-823 cells also caused cultured HUVECs to release VWF (**Figure 4C**), indicating that this endothelial cell-activating activity is common among gcEVs from multiple clones of human gastric cancer cells. PKH-26-labeled gcEVs adhered to cultured HUVECs (**Figure 4D**), which were stained extensively for VWF in intracellular granules (i.e., Weibel-Palade bodies) and on the surface, where string-like VWF structures were visible. GcEVs increased the permeability of cultured HUVECs detected by the reduction of transendothelial electrical resistance (**Figure 4E & 4F).** Similar results were also obtained using EVs from AGS and BGC-823 cells **(Supplemental Figure S7**). The effect of gcEVs on endothelial permeability was enhanced when hyperadhesive rVWF was added and reversed by the VWF-blocking antibody. However, treating cells with hyperadhesive VWF alone did not increase endothelial permeability (**Figure 4E & 4F**). This observation was further validated by detecting VWF^+^gcEV-induced the endothelial leakage of FITC dextran (**Supplemental Figure S8**). PKH26^+^ gcEVs bound to cultured HUVECs, and the adhesion was enhanced in the presence of rVWF and reduced by the anti-VWF antibody (**Figure 4G**). Some of adherent gcEVs were endocytosed by endothelial cells, as shown on the Z-axis of confocal microscopy (**Figure 4H**).

**Figure 4:**
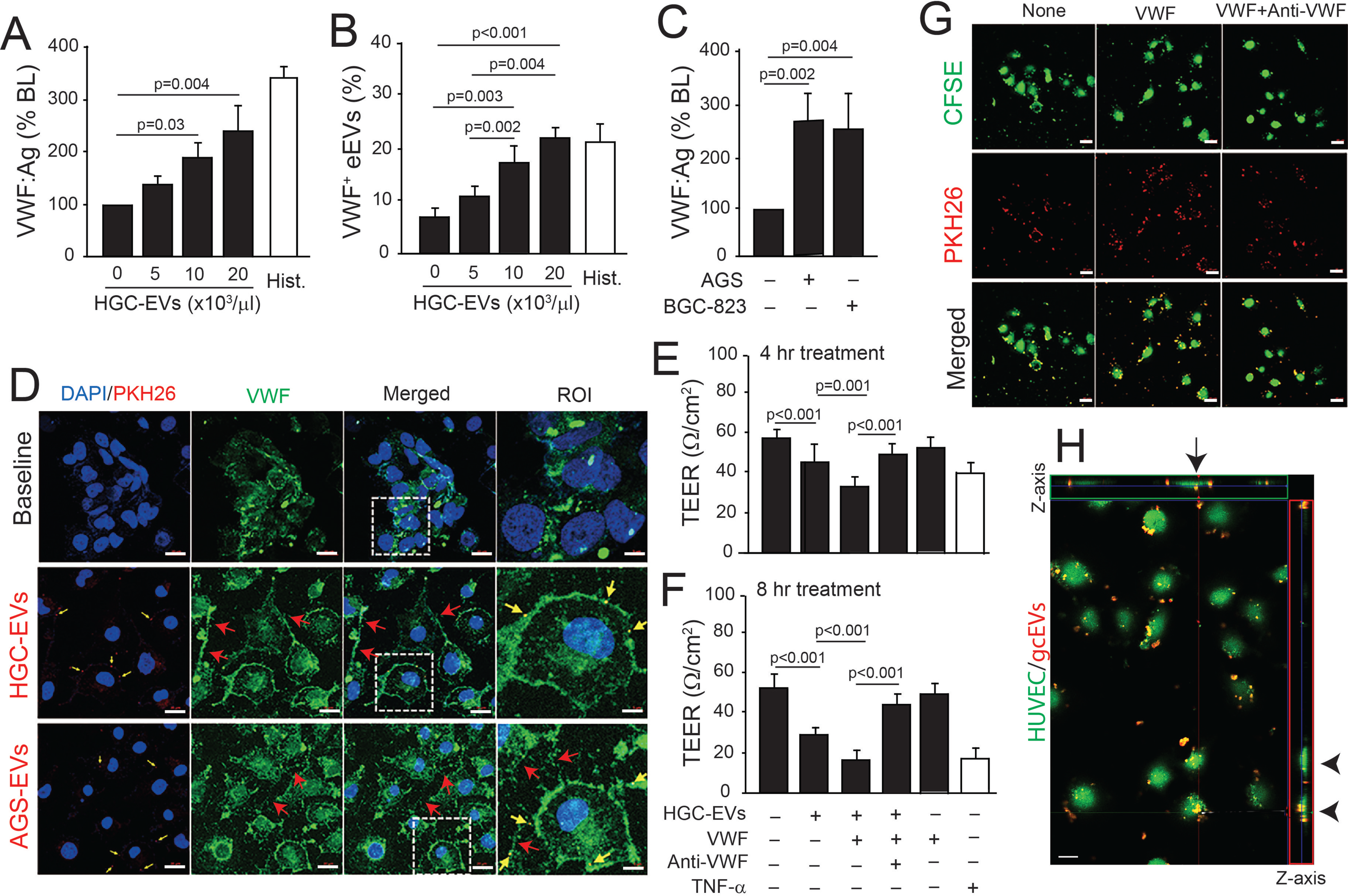
VWF enhanced gcEV-induced endothelial permeability. HGC-27-derived EVs (HGC-EVs) activated endothelial cells to release VWF (**A**) and endothelial VWF^+^/CD62E^+^ EVs (**B**) in a dose-dependent manner. Histamine-stimulated ECs as control (n=12/group, one-way ANVOA). (**C**) GcEVs from AGS and BGC-823 cells also activated ECs to release VWF (n=3/group, one-way ANOVA). (**D**) PKH-26-labeled HGC-EVs or AGS-EVs bound HUVECs (yellow arrow) and induced ECs to secrete VWF that formed string-like structures (red arrow) on the surface of ECs (DAPI for nuclear stain, bar=20 μm). The last column is images enlarged from the dotted line-defined area in the merged column to better show EV adhesion (representative from 5 independent experiments, bar=5 μm). (**E & F**) HGC-EV-induced endothelial permeability as defined by reduced TEER measured at 4 and 8 hrs, respectively, after gcEV treatment. TNF-α-stimulated cells served as control. The effect of HGC-EVs was enhanced by hyperadhesive VWF and reduced by the VWF-blocking antibody (n=6/group, one-way ANOVA). (**G**) Confocal microscopic images of HUVECs treated with HGC-EVs (2,000/μL) in the presence of hyperadhesive VWF alone or together with a VWF-blocking antibody. (**H**) A Z-axis image shows gcEVs were either adherent to the surface (arrow) or endocytosed (arrow head) by ECs (bar=20 μm). Panels G & H are representative for 5 independent experiments.

In addition to increasing endothelial permeability, human gcEVs also promoted the adhesion (**Figure 5A & 5B**) and transendothelial migration (**Figure 5C** & **Supplemental Figure S9**) of HGC-27 and AGS cells. These effects of gcEVs were enhanced in the presence of rVWF and reversed by the anti-VWF antibody (**Figure 5A-5C**). Together, these in vitro data validated the finding made in mice that (1) gcEVs activated endothelial cells to increase their permeability and to release hyperadhesive VWF and (2) hyperadhesive VWF acted together with gcEVs but not on its own to promote the adhesion and transendothelial migration of gastric cancer cells.

**Figure 5.**
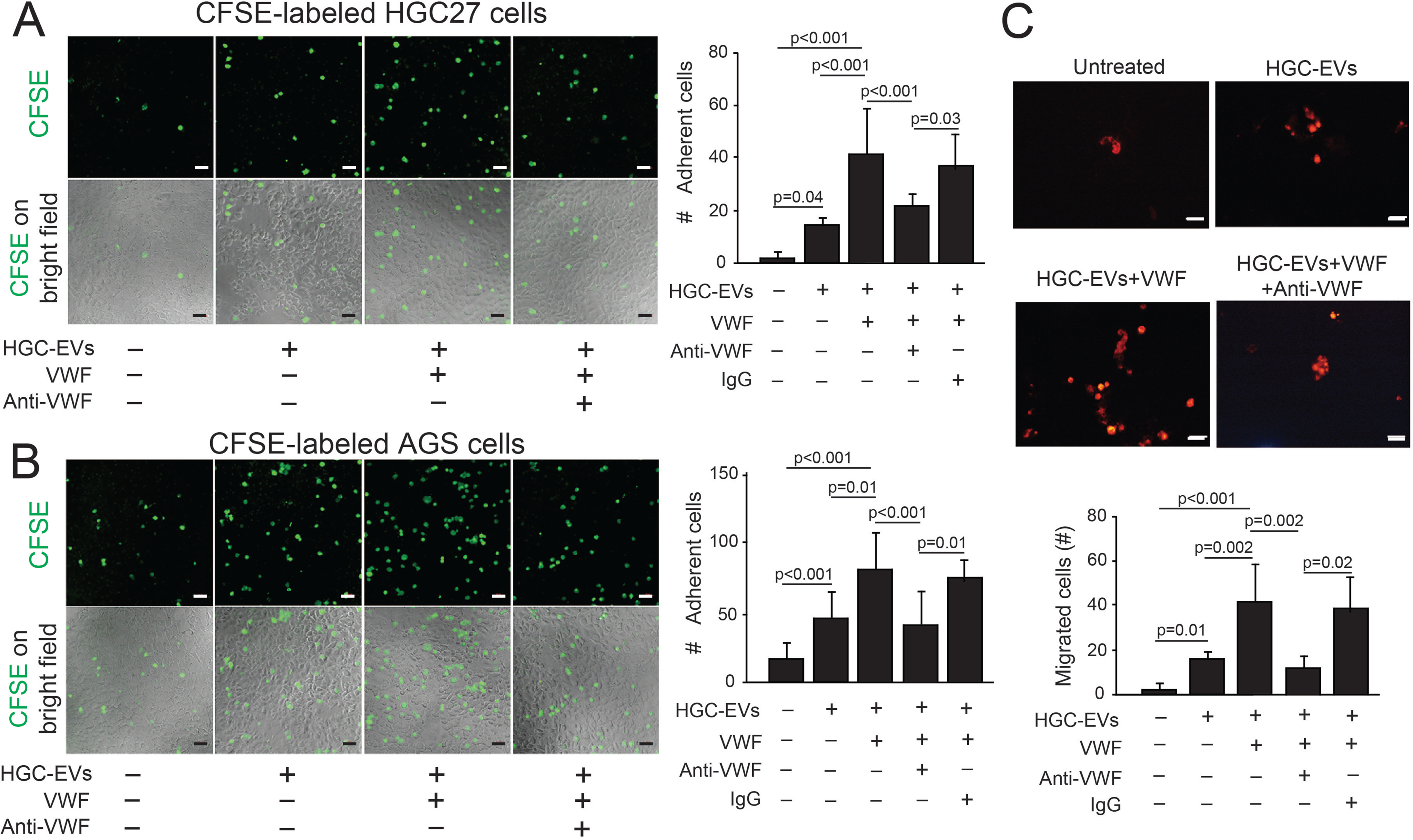
Hyperadhesive VWF enhanced the gcEV-induced adhesion of GC cells to ECs. The adhesion of CFSE-labeled human HGC-27 (**A**) and AGS cells (**B**) to the surface of ECs, showing that gcEVs increase the adhesion of HGC-27 and AGS cells and VWF further enhances the adhesion; this VWF-enhanced adhesion was blocked by the VWF antibody. Cells were seeded at 2,000/μL. Left panels: representative images from multiple experiments (bar=50 μm). Right panels: summaries of 6 independent experiments per treatment (one-way ANOVA). (**C**) Hyperadhesive VWF promoted the transendothelial transmigration of HGC-27 cells (seeding density: 5,000/μL), and the effect was blocked by the anti-VWF antibody. Left panels: representative images from multiple experiments (bar=50 μm). Right panels: summaries of 5 independent experiments per treatment (one-way ANOVA).

### VWF and gcEVs in patients with gastric cancer

The data described in Figures 1-5 demonstrate synergistic actions between hyperadhesive VWF and gcEVs in which VWF binds to gcEVs, likely through a VWF-integrin interaction.^17^ The surface-bound VWF mediates gcEV interaction with endothelial cells. These results suggest that gcEVs are released into the circulation after they interact with VWF in the tumor tissue, where vasculature is often denser and poorly developed and thus prone to leakage. To further investigate this possibility, we examined data from 380 patients diagnosed with gastric cancer and 37 patients with normal stomach tissue from the TCGA (The Cancer Genome Atlas) database^46^ for the expression of VWF in cancer and surrounding non-cancerous tissues. Analyzing data from well-established and characterized databases allowed us to study links among VWF, endothelial cells, and gastric cancer in an unbiased fashion. **Figure 6A** shows that levels of the VWF mRNA were significantly higher in STAD tissue than in noncancerous tissue. The findings were further verified by examining paired tissue samples from gastric cancer masses and adjacent normal tissue in 27 patients (**Figure 6B**) and by analyzing data from the GSE54129 and GSE19826 databases^47,48^ (**Figure 6C & 6D**). Levels of VWF mRNA expression varied significantly among patients with stage I to Ⅳ of STAD, but the highest levels were found in patients with the most advanced cancers (stage III-IV, **Supplemental Figure S10A**). As a result, the overall survival probability of patients with high levels of VWF mRNA was significantly lower than that patients with low levels of VWF mRNA (*p*=0.0153) during a follow-up period of more than 10 years (**Supplemental Figure S10B**). When the immunohistochemistry data from the Human Protein Atlas (HPA) database were examined for differential VWF expression in STAD and normal tissues, VWF was stained at a medial or strong intensity in the vascular endothelium and at a very low density in cancer cells (**Supplemental Figure S11**).

**Figure 6:**
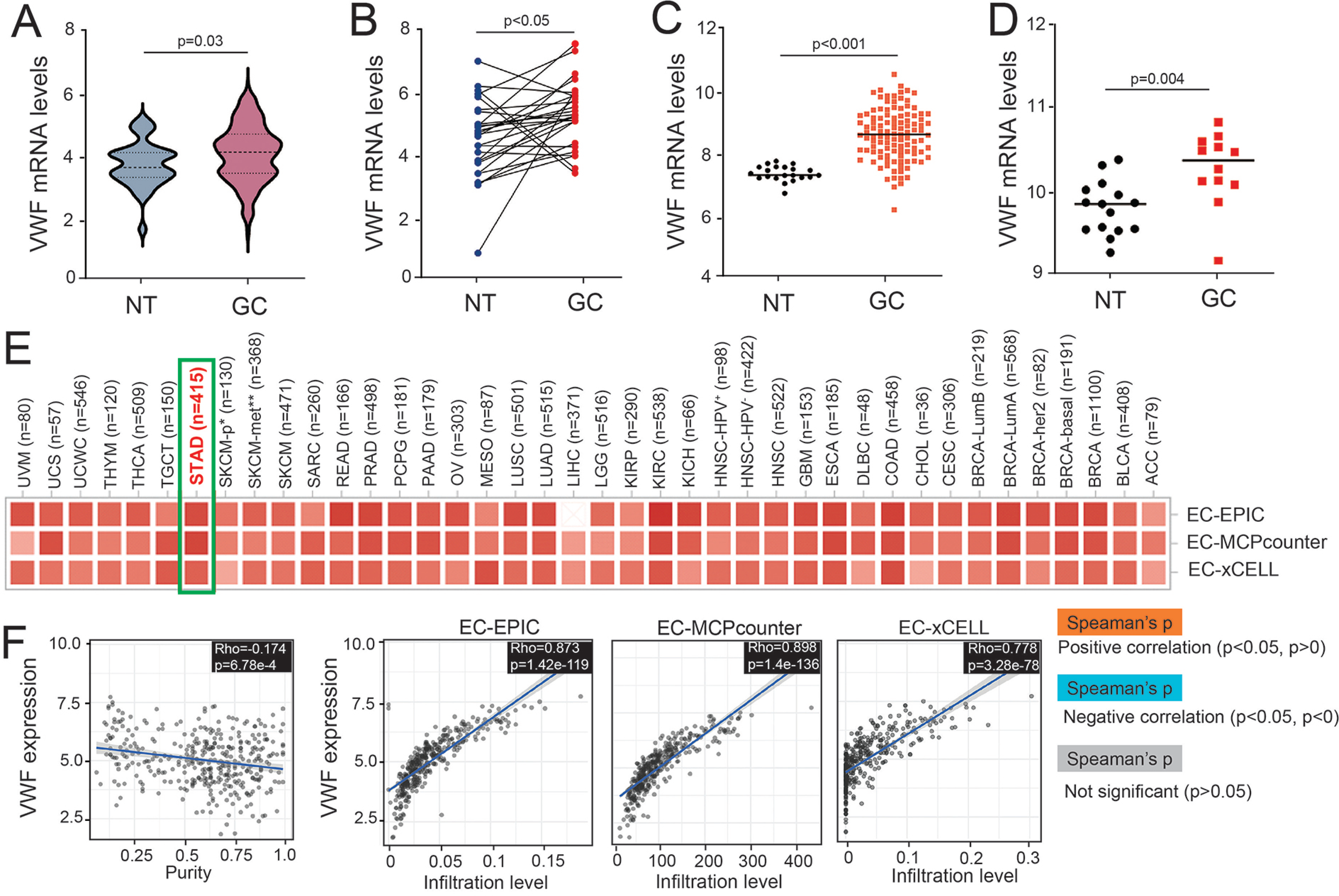
Bioinformatic analyses of VWF expression in gastric cancer tissue. (**A**) Relative levels of VWF mRNA in tissue samples from 380 GC patients and normal tissues from 37 subjects (data presented as log2, TPM+1, *t*-test). (**B**) Data validation using tissue samples from a second cohort of 27 paired samples in the STAD database from the TCGA project (independent samples t-test). The expression of VWF mRNA in gastric cancer patients and healthy controls from the GSE54129 (**C**) and GSE19826 database (**D**) was analyzed using independent samples *t*-test. (**E**) Correlation between VWF expression and endothelial cells. (**F**) Significant correlation between VWF expression and the expression of endothelial genes (log2 TPM) in STAD database through TIMER2 defined by three different algorithms (EPIC, MCP-counter, and xCell).

These data suggest that VWF was overexpressed in the endothelial cells of cancer tissues. This notion was further examined using multiple algorithms to delineate the relationship between the gene expression of endothelial cells and VWF using information from the STAD database (**Figure 6E**). The scatterplot data produced using the EPIC, MCP-counter, and xCell algorithms show that VWF expression in STAD was correlated with the level of endothelial cells (**Figure 6F**), suggesting that (1) enhanced VWF expression in cancer tissue was primarily derived from endothelial cells and (2) patients with high VWF expression in gastric cancer tissue had worse prognosis.

## DISCUSSION

We performed in vitro experiments and studied mouse models to investigate the synergistic action of VWF, gcEVs, and endothelial cells in promoting the pulmonary metastasis of gastric cancer. We also analyzed existing clinical databases to link VWF and endothelial cells to gastric cancer development. The observations made in this study are consistent with our recent clinical report that plasma samples from gastric cancer patients contain elevated levels of VWF-bound gcEVs, which associate with histopathology and clinical characteristics of the patients with more advanced cancer.^13^ This study identified novel synergies between VWF-bound gcEVs through the following novel observations. First, VWF antigen has been widely reported to be elevated in the circulation and is associated with poor clinical outcomes of gastric cancer^49^ and other types of cancers.^50,51^ However, it is not known whether VWF plays a causal role in cancer metastasis or merely serves as a marker of endothelial injury. Our study shows that VWF also became intrinsically hyperadhesive in cancer-bearing mice, especially those also infused with gcEVs (**Figure 2**), consistent with our clinical observation.^13^ VWF becomes hyperadhesive because: (1) Gastric cancer cells synthesize and release VWF;^17^ (2) VWF expression was enhanced in the endothelial cells of gastric cancer tissues (**Figure 6**), and (3) cancer-associated systemic inflammation stimulates VWF release from endothelial cells of non-cancerous tissue.^52-54^

More importantly, this elevated VWF release is not balanced by the corresponding release of ADAMTS-13,^55-57^ resulting in a kinetic deficiency of ADAMTS-13 and resultant VWF hyperadhesive activity. VWF may also undergo biochemical and conformational changes in the oxidative stress environment associated with cancer:^58,59^ (1) VWF becomes resistant to ADAMTS-13 when the methionine residue 1606 at the cleavage site is oxidized.^60^ (2) Methionine oxidation makes ADAMTS-13 less active in cleaving VWF.^61^ (3) Oxidation on the cysteine residues of VWF facilitates intermultimer disulfide bonding to form fibrillary structures^62^ that are less cleavable. (4) The A1 and A2 domains form a complex in circulating VWF multimers to make them inactive in binding plateletss and less prone to cleavage. This inhibitory complex is formed when the two vicinal cysteine residues in the A2 domain are in thiol forms but is disrupted when they are oxidized to form a disulfide bond,^63^ exposing the A1 domain for binding and activating platelets, but not facilitating sufficient cleavage because of methionine oxidation.^60^ We have detected this A1-A2 disassociation in patients supported by left ventricular assist devices^64^ and in mice subjected to severe traumatic brain injury.^29,32^ In both conditions, VWF is hyperadhesive. VWF in patients with gastric cancer may have undergone similar oxidative modifications and conformational changes, making them hyperadhesive.

Second, when given together with gcEVs but not alone, hyperadheisve VWF promotes the pulmonary metastasis of gastric cancer, which is reversed by ADAMTS-13 (**Figure 3**). An anti-VWF antibody also prevents gcEVs from adhering and activating endothelial cells to increase permeability (**Figure 4**). In this synergistic action, VWF serves as a coupling factor linking gcEVs to endothelial cells, where gcEVs activate endothelial cells and reduce the expression of the junction proteins CD31 and cadherin to promote endothelial permeability through membrane remodeling by the GTPase dynamin,^12,65^ thus increasing the transmigration of gastric cancer cells. This VWF-gcEV interaction can be mediated by the RGD sequence in the VWF C domain and integrin on gastric cancer cells,^17^ and VWF on gcEVs can then bind endothelial cells^66^ through CD62p^67^ and/or the integrin α_v_β_3_.^68,69^ The coupling effect depends on the multimeric structure of VWF with larger multimers being most effective. This explains why ADAMTS-13 reduced the pulmonary metastasis of gastric cancer (**Figure 3**), as it selectively cleaves the large and endothelial cell-bound VWF.^70^

Third, the pulmonary metastasis of gastric cancer is reduced by ADAMTS-13, which cleaves VWF to reduce its adhesive activity^71^ or by lactadherin, which removes PS-expressing EVs from the circulation^12,30^ (**Figures 1 & 3**). By binding to PS, lactadherin may also reduce cancer-associated hypercoagulable and prothrombotic states.^72,73^ While both ADAMTS-13 and anti-VWF antibody are capable of reducing gcEV-induced endothelial permeability (**Figure 5**) and gastric cancer metastasis,^17^ the former may be more targeted because ADAMTS-13 preferentially cleaves membrane-anchored and hyperadhesive VWF over cleaving the circulating plasma type of VWF.^70,74^ Although not specifically examined, platelets, which can be activated by hyperadhesive VWF, are widely recognized for their role in cancer growth and metastasis, as discussed in a recent review.^75^

In summary, we show that hyperadhesive VWF synergizes with gcEVs to induce endothelial permeability and promote the plumonary metastasis of gastric cancer. This synergistic action is likely a self-propagating process in which apoptotic cancer cells release EVs that becomes VWF-bound when they are released from the poorly developed and “leaky” vessels in the cancer tissue. When in the circulation, VWF enhances the EV-induced activation of endothelial cells to release more hyperadhesive VWF, which can bind gcEVs to promote metastasis and to activate platelets to release yet more VWF and PS^+^EVs to propage a prothrombotic state. Reducing PS^+^EVs and blocking hyperadhesive VWF may therefore be new strategies to reduce the metastasis of gastric cancer.

## Acknowledgement

We are very grateful for having the access to the TCGA, GEO, TIMER2, HPA database for specific analyzes. This research was supported by grants from the National Natural Science Foundation of China 81871919 and 81672399 (ML), the Natural Science Foundation of Gansu Province 22JR5RA494 (CYW) and 22JR5RA501 (MW), the Fundamental Research Funds for the Central Universities lzujbky-2017-136 (CYW), the Excellent graduate student project 2022CXZX-131 (CYZ).

## Author contributions

Chen-yu Wang designed and performed experiments, analyzed data, and wrote manuscript; Min Wang designed and performed experiments, analyzed data; Wei Cai collected the data and performed experiments; Chan-yuan Zhao and Quan Zhou performed mouse experiments; Xiao-yu Zhang, Feng-xia Wang, and Chen-li Zhang performed experiments; Yun Dang and Ai-jun Yang completed database data analysis; Jing-fei Dong and Min Li developed hypotheses, designed the study, analyzed data, and wrote manuscript. All authors read and approved the final manuscript.

## Conflict of interest statement

The authors claim no conflict of interest relevant to this study.

## Notes

### Competing Interest Statement

The authors have declared no competing interest.

## REFERENCES

1. Chiang TH, Chang WJ, Chen SL, et al. Mass eradication of Helicobacter pylori to reduce gastric cancer incidence and mortality: a long-term cohort study on Matsu Islands. Gut. 2021;70(2):243–250.

2. Sung H, Ferlay J, Siegel RL, et al. Global Cancer Statistics 2020: GLOBOCAN Estimates of Incidence and Mortality Worldwide for 36 Cancers in 185 Countries. CA Cancer J Clin. 2021;71(3):209-249.

3. Qiu H, Cao S, Xu R. Cancer incidence, mortality, and burden in China: a time-trend analysis and comparison with the United States and United Kingdom based on the global epidemiological data released in 2020. Cancer Commun (Lond). 2021;41(10):1037–1048.

4. Entenberg D, Voiculescu S, Guo P, et al. A permanent window for the murine lung enables high-resolution imaging of cancer metastasis. Nat Methods. 2018;15(1):73–80.

5. Aran D, Camarda R, Odegaard J, et al. Comprehensive analysis of normal adjacent to tumor transcriptomes. Nat Commun. 2017;8(1):1077.

6. Vlodavsky I, Gross-Cohen M, Weissmann M, Ilan N, Sanderson RD. Opposing Functions of Heparanase-1 and Heparanase-2 in Cancer Progression. Trends Biochem Sci. 2018;43(1):18–31.

7. Abd Elmageed ZY, Yang Y, Thomas R, et al. Neoplastic reprogramming of patient-derived adipose stem cells by prostate cancer cell-associated exosomes. Stem Cells. 2014;32(4):983–997.

8. Peinado H, Zhang H, Matei IR, et al. Pre-metastatic niches: organ-specific homes for metastases. Nature Reviews Cancer. 2017;17(5):302–317.

9. Wang CC, Tseng CC, Hsiao CC, et al. Circulating endothelial-derived activated microparticle: a useful biomarker for predicting one-year mortality in patients with advanced non-small cell lung cancer. Biomed Res Int. 2014;2014:173401.

10. Galindo-Hernandez O, Villegas-Comonfort S, Candanedo F, et al. Elevated concentration of microvesicles isolated from peripheral blood in breast cancer patients. Arch Med Res. 2013;44(3):208–214.

11. Wang W, Li H, Zhou Y, Jie S. Peripheral blood microvesicles are potential biomarkers for hepatocellular carcinoma. Cancer Biomark. 2013;13(5):351–357.

12. Wang M, Cai W, Yang AJ, et al. Gastric cancer cell-derived extracellular vesicles disrupt endothelial integrity and promote metastasis. Cancer Lett. 2022;545:215827.

13. Wei Cai, Min Wang, Chen-yu Wang, et al. Extracellular Vesicles, Hyperadhesive von Willebrand Factor, and Outcomes of Gastric Cancer: a Clinical Observational Study. Medical oncology. 2023;40:140.

14. Shao H, Im H, Castro CM, Breakefield X, Weissleder R, Lee H. New Technologies for Analysis of Extracellular Vesicles. Chem Rev. 2018;118(4):1917–1950.

15. D’Acunzo P, Kim Y, Ungania JM, Pérez-González R, Goulbourne CN, Levy E. Isolation of mitochondria-derived mitovesicles and subpopulations of microvesicles and exosomes from brain tissues. Nat Protoc. 2022;17(11):2517–2549.

16. O’Brien K, Breyne K, Ughetto S, Laurent LC, Breakefield XO. RNA delivery by extracellular vesicles in mammalian cells and its applications. Nat Rev Mol Cell Biol. 2020;21(10):585–606.

17. Yang AJ, Wang M, Wang Y, et al. Cancer cell-derived von Willebrand factor enhanced metastasis of gastric adenocarcinoma. Oncogenesis. 2018;7(1):12.

18. Lenting PJ, Christophe OD, Denis CV. von Willebrand factor biosynthesis, secretion, and clearance: connecting the far ends. Blood. 2015;125(13):2019–2028.

19. Levy GG, Nichols WC, Lian EC, et al. Mutations in a member of the ADAMTS gene family cause thrombotic thrombocytopenic purpura. Nature. 2001;413(6855):488-494.

20. Zheng X, Chung D, Takayama TK, Majerus EM, Sadler JE, Fujikawa K. Structure of von Willebrand factor-cleaving protease (ADAMTS13), a metalloprotease involved in thrombotic thrombocytopenic purpura. J Biol Chem. 2001;276(44):41059–41063.

21. Dong JF, Moake JL, Nolasco L, et al. ADAMTS-13 rapidly cleaves newly secreted ultralarge von Willebrand factor multimers on the endothelial surface under flowing conditions. Blood. 2002;100(12):4033–4039.

22. Chen J, Chung DW. Inflammation, von Willebrand factor, and ADAMTS13. Blood. 2018;132(2):141–147.

23. Sadler JE. Pathophysiology of thrombotic thrombocytopenic purpura. Blood. 2017;130(10):1181–1188.

24. Goh CY, Patmore S, Smolenski A, et al. The role of von Willebrand factor in breast cancer metastasis. Transl Oncol. 2021;14(4):101033.

25. Gu J, Qi Y, Lu Y, et al. Lung adenocarcinoma-derived vWF promotes tumor metastasis by regulating PHKG1-mediated glycogen metabolism. Cancer Sci. 2022;113(4):1362–1376.

26. Patmore S, Dhami SPS, O’Sullivan JM. Von Willebrand factor and cancer; metastasis and coagulopathies. J Thromb Haemost. 2020;18(10):2444–2456.

27. Cheung EC, DeNicola GM, Nixon C, et al. Dynamic ROS Control by TIGAR Regulates the Initiation and Progression of Pancreatic Cancer. Cancer Cell. 2020;37(2):168–182 e164.

28. Han Y, Xiao J, Falls E, Zheng XL. A shear-based assay for assessing plasma ADAMTS13 activity and inhibitors in patients with thrombotic thrombocytopenic purpura. Transfusion. 2011;51(7):1580–1591.

29. Yang M, Houck KL, Dong X, et al. Hyperadhesive von Willebrand Factor Promotes Extracellular Vesicle-Induced Angiogenesis: Implication for LVAD-Induced Bleeding. JACC Basic Transl Sci. 2022;7(3):247–261.

30. Zhou Y, Cai W, Zhao Z, et al. Lactadherin promotes microvesicle clearance to prevent coagulopathy and improves survival of severe TBI mice. Blood. 2018;131(5):563–572.

31. Groeneveld D, Cline-Fedewa H, Baker KS, et al. Von Willebrand factor delays liver repair after acetaminophen-induced acute liver injury in mice. J Hepatol. 2020;72(1):146–155.

32. Wu Y, Liu W, Zhou Y, et al. von Willebrand factor enhances microvesicle-induced vascular leakage and coagulopathy in mice with traumatic brain injury. Blood. 2018;132(10):1075–1084.

33. Han C, Wang C, Chen Y, et al. Placenta-derived extracellular vesicles induce preeclampsia in mouse models. Haematologica. 2020;105(6):1686–1694.

34. Bernardo A, Ball C, Nolasco L, Moake JF, Dong JF. Effects of inflammatory cytokines on the release and cleavage of the endothelial cell-derived ultralarge von Willebrand factor multimers under flow. Blood. 2004;104(1):100–106.

35. Paraiso KH, Das Thakur M, Fang B, et al. Ligand-independent EPHA2 signaling drives the adoption of a targeted therapy-mediated metastatic melanoma phenotype. Cancer Discov. 2015;5(3):264–273.

36. Albrengues J, Shields MA, Ng D, et al. Neutrophil extracellular traps produced during inflammation awaken dormant cancer cells in mice. Science. 2018;361(6409).

37. Reichert M, Bakir B, Moreira L, et al. Regulation of Epithelial Plasticity Determines Metastatic Organotropism in Pancreatic Cancer. Dev Cell. 2018;45(6):696–711 e698.

38. Nagorcka-Smith P, Bolton KA, Dam J, et al. The impact of coalition characteristics on outcomes in community-based initiatives targeting the social determinants of health: a systematic review. BMC Public Health. 2022;22(1):1358.

39. Wang G, Lu X, Dey P, et al. Targeting YAP-Dependent MDSC Infiltration Impairs Tumor Progression. Cancer Discov. 2016;6(1):80–95.

40. Ding Z, Wu CJ, Chu GC, et al. SMAD4-dependent barrier constrains prostate cancer growth and metastatic progression. Nature. 2011;470(7333):269-273.

41. Li T, Fu J, Zeng Z, et al. TIMER2.0 for analysis of tumor-infiltrating immune cells. Nucleic Acids Res. 2020;48(W1):W509–W514.

42. Becht E, Giraldo NA, Lacroix L, et al. Estimating the population abundance of tissue-infiltrating immune and stromal cell populations using gene expression. Genome Biol. 2016;17(1):218.

43. Aran D, Hu Z, Butte AJ. xCell: digitally portraying the tissue cellular heterogeneity landscape. Genome Biol. 2017;18(1):220.

44. Racle J, de Jonge K, Baumgaertner P, Speiser DE, Gfeller D. Simultaneous enumeration of cancer and immune cell types from bulk tumor gene expression data. Elife. 2017;13(6):e26476.

45. Greten FR, Grivennikov SI. Inflammation and Cancer: Triggers, Mechanisms, and Consequences. Immunity. 2019;51(1):27–41.

46. Cancer Genome Atlas Research N. Comprehensive molecular characterization of gastric adenocarcinoma. Nature. 2014;513(7517):202-209.

47. Liu AG, Zhong JC, Chen G, et al. Upregulated expression of SAC3D1 is associated with progression in gastric cancer. Int J Oncol. 2020;57(1):122–138.

48. Shen H, Wang L, Chen Q, et al. The prognostic value of COL3A1/FBN1/COL5A2/SPARC-mir-29a-3p-H19 associated ceRNA network in Gastric Cancer through bioinformatic exploration. J Cancer. 2020;11(17):4933–4946.

49. Yang X, Sun HJ, Li ZR, et al. Gastric cancer-associated enhancement of von Willebrand factor is regulated by vascular endothelial growth factor and related to disease severity. BMC Cancer. 2015;15:80.

50. Marfia G, Navone SE, Fanizzi C, et al. Prognostic value of preoperative von Willebrand factor plasma levels in patients with Glioblastoma. Cancer Med. 2016;5(8):1783–1790.

51. Wei-Shu Wang J-KL, Tzu-Chen Lin, Tzeon-Jye Chiou, Jin-Hwang Liu, Chueh-Chuan Yen, Po-Min Chen. Plasma von Willebrand factor level as a prognostic indicator of patients with metastatic colorectal carcinoma. World J Gastroenterol. 2005;11(14):2166–2170.

52. Junmei Chen DWC. Inflammation, von Willebrand factor, and ADAMTS13. Blood. 2018;132(2):141–147.

53. Mantovani A, Allavena P, Sica A, Balkwill F. Cancer-related inflammation. Nature. 2008;454(7203):436-444.

54. Strozyk EA, Desch A, Poeppelmann B, et al. Melanoma-derived IL-1 converts vascular endothelium to a proinflammatory and procoagulatory phenotype via NFκB activation. Exp Dermatol. 2014;23(9):670–676.

55. Guo R, Yang J, Liu X, Wu J, Chen Y. Increased von Willebrand factor over decreased ADAMTS-13 activity is associated with poor prognosis in patients with advanced non-small-cell lung cancer. J Clin Lab Anal. 2018;32(1):e22219.

56. Takaya H, Namisaki T, Moriya K, et al. Association between ADAMTS13 activity-VWF antigen imbalance and the therapeutic effect of HAIC in patients with hepatocellular carcinoma. World J Gastroenterol. 2020;26(45):7232–7241.

57. Obermeier HL, Riedl J, Ay C, et al. The role of ADAMTS-13 and von Willebrand factor in cancer patients: Results from the Vienna Cancer and Thrombosis Study. Res Pract Thromb Haemost. 2019;3(3):503–514.

58. Sonowal H, Shukla K, Kota S, Saxena A, Ramana KV. Vialinin A, an Edible Mushroom-Derived p-Terphenyl Antioxidant, Prevents VEGF-Induced Neovascularization In Vitro and In Vivo. Oxid Med Cell Longev. 2018;2018:1052102.

59. Morry J, Ngamcherdtrakul W, Yantasee W. Oxidative stress in cancer and fibrosis: Opportunity for therapeutic intervention with antioxidant compounds, enzymes, and nanoparticles. Redox Biol. 2017;11:240–253.

60. Chen J, Fu X, Wang Y, et al. Oxidative modification of von Willebrand factor by neutrophil oxidants inhibits its cleavage by ADAMTS13. Blood. 2010;115(3):706–712.

61. Wang Y, Chen J, Ling M, Lopez JA, Chung DW, Fu X. Hypochlorous acid generated by neutrophils inactivates ADAMTS13: an oxidative mechanism for regulating ADAMTS13 proteolytic activity during inflammation. J Biol Chem. 2015;290(3):1422–1431.

62. Choi H, Aboulfatova K, Pownall HJ, Cook R, Dong JF. Shear-induced disulfide bond formation regulates adhesion activity of von Willebrand factor. J Biol Chem. 2007;282(49):35604–35611.

63. Diego Butera FP, Lining Ju, Kristina M. Cook, Heng Woon, Camilo Aponte-Santamaría, Elizabeth Gardiner, Amanda K. Davis, Deirdre A. Murphy, Agnieszka Bronowska, Brenda M. Luken, Carsten Baldauf, Shaun Jackson, Robert Andrews, Frauke Gräter, Philip J. Hogg. Autoregulation of von Willebrand factor function by a disulfide bond switch. SCIENCE ADVANCES. 2018;4:eaaq1477.

64. Xin Xu, Chenyu Wang, Yingang Wu, et al. Conformation-dependent blockage of activated VWF improves outcomes of traumatic brain injury in mice. Blood. 2021;137(4):544–555.

65. Ferguson SM, De Camilli P. Dynamin, a membrane-remodelling GTPase. Nat Rev Mol Cell Biol. 2012;13(2):75–88.

66. Tian Y, Salsbery B, Wang M, et al. Brain-derived microparticles induce systemic coagulation in a murine model of traumatic brain injury. Blood. 2015;125(13):2151–2159.

67. Padilla A, Moake JL, Bernardo A, et al. P-selectin anchors newly released ultralarge von Willebrand factor multimers to the endothelial cell surface. Blood. 2004;103(6):2150–2156.

68. Fuentes P, Sese M, Guijarro PJ, et al. ITGB3-mediated uptake of small extracellular vesicles facilitates intercellular communication in breast cancer cells. Nat Commun. 2020;11(1):4261.

69. Huang J, Roth R, Heuser JE, Sadler JE. Integrin alpha(v)beta(3) on human endothelial cells binds von Willebrand factor strings under fluid shear stress. Blood. 2009;113(7):1589–1597.

70. Dong JF, Moake JL, Nolasco L, et al. ADAMTS-13 rapidly cleaves newly secreted ultralarge von Willebrand factor multimers on the endothelial surface under flowing conditions. Blood. 2002;100(12):4033–4039.

71. Zander CB, Cao W, Zheng XL. ADAMTS13 and von Willebrand factor interactions. Curr Opin Hematol. 2015;22(5):452–459.

72. Hisada Y, Mackman N. Cancer-associated pathways and biomarkers of venous thrombosis. Blood. 2017;130(13):1499–1506.

73. Geddings JE, Mackman N. Tumor-derived tissue factor-positive microparticles and venous thrombosis in cancer patients. Blood. 2013;122(11):1873–1880.

74. Kenji Nishio PJA, X. Long Zheng, and J. Evan Sadler. Binding of platelet glycoprotein Ib to von Willebrand factor domain A1 stimulates the cleavage of the adjacent domain A2 by ADAMTS13. The Proceedings of the National Academy of Sciences. 2004;101(29):10578–10583.

75. Haemmerle M, Stone RL, Menter DG, Afshar-Kharghan V, Sood AK. The Platelet Lifeline to Cancer: Challenges and Opportunities. Cancer Cell. 2018;33(6):965–983.

